# Heterogeneity of RNA editing in mesothelioma and how RNA editing enzyme ADAR2 affects mesothelioma cell growth, response to chemotherapy and tumor microenvironment

**DOI:** 10.1101/2022.07.12.499727

**Authors:** Ananya Hariharan, Weihong Qi, Hubert Rehrauer, Licun Wu, Manuel Ronner, Martin Wipplinger, Jelena Kresoja-Rakic, Suna Sun, Lucia Oton-Gonzalez, Marika Sculco, Véronique Serre-Beinier, Clément Meiller, Christophe Blanquart, Jean-François Fonteneau, Bart Vrugt, Jan Hendrik Rüschoff, Isabelle Opitz, Didier Jean, Marc de Perrot, Emanuela Felley-Bosco

## Abstract

We previously observed increased levels of adenosine-deaminase-acting-on-dsRNA (Adar)-dependent RNA editing during mesothelioma development in mice exposed to asbestos. The aim of this study was to characterize and assess the role of ADAR-dependent RNA editing in mesothelioma. Tumors and mesothelioma primary cultures have higher ADAR-mediated RNA editing compared to mesothelial cells. Unsupervised clustering of editing in different genomic regions revealed heterogeneity between tumor samples as well as mesothelioma primary cultures. ADAR2 expression levels are higher in BRCA1-associated protein 1 wild-type tumors, with corresponding changes in RNA editing in transcripts and 3’UTR. ADAR2 knockdown and rescue models indicated a role in cell proliferation, altered cell cycle, increased sensitivity to antifolate treatment and type-1 interferon signaling upregulation, leading to changes in the microenvironment *in vivo*. Our data indicate that RNA editing contributes to mesothelioma heterogeneity and highlights an important role of ADAR2 not only in growth regulation in mesothelioma but also chemotherapy response, in addition to regulating inflammatory response downstream of sensing nucleic acid structures.

## Background

Pleural mesothelioma (PM) is the most common primary neoplasm arising from the mesothelial cell layer (1). It is a rapidly fatal and highly resilient (2) tumor primarily associated with previous exposure to asbestos fibers. Although the use of asbestos has been banned in most western countries, developing nations continue to use asbestos (3). This indicates that the incidence of mesothelioma will continue to rise in the years to come.

Several recent high throughput studies have documented the heterogeneity of PM, at the genetic and the transcriptomic level (4-6). Transcriptome heterogeneity can be further enhanced by RNA editing. A-to-I RNA editing is the most prevalent form of RNA editing in higher eukaryotes. It is catalyzed by enzymes known as adenosine deaminases acting on double-stranded RNA (dsRNA) or ADARs. More than 85% of human primary transcripts undergo RNA editing. A large proportion of these editing sites are within repetitive elements in untranslated regions (UTR) of transcripts, such as Alu elements, which have the ability to form secondary dsRNA structures (7). ADAR activity results in the hydrolytic deamination of adenosine to form inosine, which is then interpreted as guanosine (8). The molecular consequences of ADAR activity largely depend on the region of RNA that is targeted. For example, editing in coding sequences can change the encoded amino acid, in introns can affect alternative splicing of transcripts, and in UTR can alter RNA stability or the translation efficiency (9). Destabilization of dsRNA structures upon ADAR activity suppresses the cellular response to dsRNA by preventing type-1 interferon (IFN) production and downregulation of apoptotic and autophagic responses (10). Interestingly, in The Cancer Genome Atlas (TCGA) study (6), type-1 IFN signaling is associated with the mutational status of BRCA1-associated protein (BAP1), a tumor suppressor gene, which is frequently inactivated in mesothelioma (reviewed in (1)), however the underlying mechanisms are not clear.

The mammalian ADAR family is comprised of three members - ADAR1, ADAR2, and ADAR3. ADAR3 is inactive and known to be expressed only in the central nervous system (11). All three ADARs contain a conserved C-terminal deaminase domain and multiple RNA binding domains.

Dysregulation of ADAR expression and ADAR-mediated RNA editing has been shown to result in various diseases including cancer (12). Both ADAR1 and ADAR2 have roles in regulating genomic stability in cancer cells. ADAR1 p110 was found to edit RNA in DNA:RNA hybrids leading to the resolution of R-loops, promoting proliferation of cancer cells (13). Similarly, dissolution of DNA:RNA hybrids by ADAR2-mediated editing rendered cancer cells resistant to genotoxic agents and decreases genomic instability (14).

In previous published work on mesothelioma development, we have found that mice chronically exposed to asbestos show a significant increase in A-to-I RNA editing even in pre-neoplastic lesions, which further increased in tumors (15, 16). *Adar1* expression increased upon asbestos-exposure in inflamed tissues, but *Adar2* expression increased only upon tumor formation (15). This was consistent with the observed upregulation of Yes-associated protein and Transcriptional co-activator with PDZ binding motif (YAP/TAZ) pathway (15) alongside decrease in ADAR2 expression upon blocking the YAP/TAZ pathway in mesothelioma cells (17). Sakata, et al. showed by gene silencing that ADAR2 but not ADAR1 inhibited TCC-MESO1 mesothelioma cells growth (18), but the mechanisms have not been thoroughly explored.

The aim of this study was to characterize the ADAR-dependent RNA editing landscape in human mesothelioma and put it into context with known BAP1 status. We established stable ADAR2 deficient cells and their rescue to identify the pathways through which ADAR2 exerts its role in cell growth, cell cycle progression, and interaction with the microenvironment. We also determined the effect of ADAR2 loss in response to standard chemotherapy.

## Materials and Methods

### A to G index computing and stratification

Mesothelioma RNA-seq reads included in the analysis were: the TCGA-Mesothelioma cohort (n = 87) downloaded from the NCBI database of Genotypes and Phenotypes (dbGaP) in 2019, under phs000178.v10.p8; the Malignant Pleural Mesothelioma cohort from the Bueno study (n=223) downloaded from the European Genome-phenome Archive (EGA) in 2020, under EGAS00001001563 (EGAD00001001915 and EGAD00001001916); the genetically characterized pleural mesothelioma primary cultures (n=64) provided by Didier Jean’s team in 2022 for which RNA-Seq was performed as described in (19); the human pluripotent stem cell derived mesothelium (n=10) downloaded from the NCBI Gene Expression Omnibus (GEO) in 2020, under GSE113090 (GSM3096389-GSM3096398). The choice of using human pluripotent stem cell derived mesothelium as control was dictated by the fact that RNA-Seq data on normal mesothelial cells or pleura are not available and RNA editing is tissue-specific (20). RNA-seq reads were pre-processed using fastp (0.20.0). The first 6 bases at the beginning of each read were deleted to remove priming bias (21) introduced during Illumina RNA-seq library preparation. Sequencing adapters and low-quality ends (averaged quality lower than 20 in sliding windows of 4 bp, moving from 5’ to 3’, and from 3’ to 5’, respectively) were trimmed. Reads longer than 48 nt were trimmed back to 48 nt, in order to achieve uniform max read length across different datasets and comparable RNA editing index values. Trimmed reads with average quality above 20 and length between 18 and 48 bp were aligned to the human reference genome (Genomic Data Commons (GDC) GRCh38.d1.vd1 Reference Sequence, https://gdc.cancer.gov/about-data/gdc-data-processing/gdc-reference-files) using STAR (2.7.8a) with 2-pass mode. PCR duplicates were marked using Picard (2.22.8).

Primary alignments were extracted using samtools (1.11) and were used for computing the A to G index and C to T index by applying the python package RNAEditingIndexer (https://github.com/a2iEditing/RNAEditingIndexer).

Variants in the aligned, duplicate marked RNA-seq reads were identified using GATK (v3.8.1.0) following RNA-seq best practices workflows. In details, mapping quality reassignment, splitting spliced aligned reads into multiple supplementary alignments and clipping mismatching overhangs were performed using “SplitNCigarReads” with options “-rf ReassignOneMappingQuality -RMQF 255 -RMQT 60 -U ALLOW_N_CIGAR_READS”. Base quality recalibration was performed using “BaseRecalibrator” with dbSNP release151_GRCh38p7 downloaded in 2018 as the true variant set. Variant calling was performed using “HaplotypeCaller” with options “-dontUseSoftClippedBases -stand_call_conf 20.0”. Called variants were filtered using “VariantFiltration” with the following options: -window 35 -cluster 3 - filterName FS -filter “FS > 30.0” -filterName QD -filter “QD < 2.0”. Variants known in dbSNP, or/and in genes encode immunoglobulins were also filtered out using SnpSift (v4.3) and bedtools (v2.29.2) respectively. Filtered variants were annotated using SnpEff (v4.3) and GDC.h38 GENCODE v22 gene annotation. Percentages of A to G changes by genomic regions in the SnpEff csv summary file were extracted for sample clustering analysis in R (v4.1). In details, pairwise euclidean distances among samples were computed based on percentage values of A to G changes by genomic regions with R function “dist”. Sample clustering based on the distance matrix was performed using the Ward’s hierarchical clustering method as implemented in “hclust”. Sample grouping was determined by cutting the resulting dendrogram tree (cuttree) into groups by specifying six clusters (k=6). To compare editing to genomic regions content in the human genome, the sum of the percentages of SnpEff calculated editing in “exons”, “transcripts”, “5’UTR”, “3’UTR” were grouped under exons.

To test for sequence motifs, a region ±5 nucleotides around the editing site was analyzed using pLogo generator (https://plogo.uconn.edu/)(22).

Gene expression values from aligned RNA-seq reads were computed using htseq-count (v1.99.2) with options “-a 10 -t exon -i gene_id -m intersection-nonempty”. For the stranded FunGeST dataset, “-s reverse” was set, while “-s no” was used for all other non-stranded datasets. FPKM and FPKM-UQ values were computed using R (v4.1), where transcript length information was downloaded from GDC (“genecode.gene.info.v22.tsv”).

Mutational status of *BAP1* was extracted from TCGA (6) and Bueno’s (5, 23) datasets. Clustering of Bueno’s dataset samples with wild-type (wt) BAP1 in two groups was included due to the fact that recent analysis (23) uncovered events masked by abundant stroma. Primary pleural mesothelioma cell lines of FunGeST series were characterized for genetic alterations in key genes of mesothelial carcinogenesis including *BAP1* using a targeted sequencing described in (24).

Analysis of *ADAR2* co-expressed genes with a positive Pearson correlation coefficient (r >0.296, q < 0.05) in cBioPortal.org mesothelioma TCGA dataset was selected for transcription factor enrichment annotation using the ENRICHR (25) online platform.

### Cell culture

Mesothelioma cell lines (Table S1) were cultured as previously described (16, 26). Cell lines were confirmed to be free of Mycoplasma on a regular basis using PCR Mycoplasma kit (MD Bioproducts).

### Protein extraction and Western blotting

Total protein extracts were obtained by lysing the cells with hot Laemmli buffer (60 mM Tris-HCl pH 6.8, 100 mM DTT, 5% glycerol, 1.7% SDS) and passed through syringes (26G). A total of 5 μg protein extract was separated on denaturing 10, 15 or 10-20% SDS-PAGE gels and proteins were transferred onto PVDF membranes (0.45 μm, Perkin Elmer, Waltham, MA) and blocked with 5% BSA/TBS-T for 2 hours at room temperature. Membranes were probed with the following primary antibodies, ADAR1 (Sigma-Aldrich Cat# HPA003890, RRID:AB_1078103), ADAR2 (Sigma-Aldrich Cat# HPA018277, RRID:AB_1844591; Santa Cruz Biotechnology Cat# sc-73409, RRID:AB_2289194), Flag-tag (Sigma-Aldrich Cat# F1804, RRID:AB_262044), DHFR (Santa Cruz Biotechnology Cat# sc-377091, RRID:AB_) TYMS (Santa Cruz Biotechnology Cat# sc-33679, RRID:AB_628355), RIG-I (Cell Signaling Technology Cat# 3743, RRID:AB_2269233), IFITM1 (Novus Cat# NBP1-77171, RRID:AB_11010388), MAVS (Mouse: Santa Cruz Biotechnology Cat# sc-365333, RRID:AB_10844335, Human: Cell Signaling Technology Cat# 3993, RRID: AB_823565), STING (Cell Signaling Technology Cat# 13647, RRID:AB_2732796), ISG15 (Santa Cruz Biotechnology Cat# sc-166755, RRID:AB_2126308) and β-actin (C4, MP Bio-medicals MP691002 RRID:AB_2335127), overnight at 4°C. Membranes were then incubated with one of the following secondary antibodies at room temperature for 1 hour: rabbit anti-mouse IgG-HRP (no. A9004) or goat anti-rabbit IgG-HRP (no. A0545), obtained from Sigma Aldrich. Signals were detected with enhanced chemiluminescence reagent (Clarity TM ECL Substrate, BioRad, Hercules, CA) using Fusion Digital Imager (Vilber Lourmat, Marne-la-Vallée, France). Quantification of bands was done using ImageJ software.

### Patient cohort

Tumor tissue was collected from 193 patients between 1999-2015 (Table S2). A subset of patients has been treated with a neoadjuvant chemotherapy consisting of cisplatin and pemetrexed, followed by surgery (P/D or EPP) at the University Hospital of Zurich. The study was approved by the Ethical Committee Zürich (KEK-ZH-2012-0094 and BASEC-No. 2020-02566), and patients either signed informed consent or waiver of consent was granted by the Ethical Committee (BASEC-No. 2020-02566). The study methodologies were conformed to the standards set by the Declaration of Helsinki. Response to chemotherapy was assessed in PET-CT Scans according to the modified RECIST (mRECIST) criteria (27).

### Tissue microarray

TMA construction was achieved as previously described (28). Immunohistochemistry was performed as previously described (29) using rabbit anti-ADAR2 (Santa Cruz Biotechnology Cat# sc-73409, RRID:AB_2289194) antibody. TMA spots with a lack of tumor tissue were excluded from the analysis, resulting in the analysis of 178 patients’ samples. Immunohistochemical evaluation of the TMAs was conducted by two independent observers in a blinded manner. The average H-score of all the cores obtained from the same tumor specimen was determined by the percentage of cells having nuclear ADAR2 positivity and scored as 0 (0%), 1 (1%–9%), 2 (10%–49%), or 3 (50% and more).

### RNA extraction, cDNA synthesis, RT-qPCR and AZIN1 editing

0.5μg of total RNA was extracted from cells using RNeasy isolation kit (QIAGEN, Cat No.74106) and reverse-transcribed using the Quantitect Reverse Transcription Kit (QIAGEN, Cat No.205311) according to the manufacturer’s instructions. Synthesized cDNA was diluted 1:60 or 1:36 (for spheroids) and used for real-time quantitative PCR (RT-qPCR). SYBR green (Thermo Fisher, Cat No.4367659) and gene specific primers were used for PCR amplification and detection on a 7500 FAST Real-Time PCR System (Applied Biosystems, Thermo Fisher Scientific). Relative mRNA levels were determined by comparing the PCR cycle thresholds between cDNA of a specific gene and beta actin for mouse or histones for human (ΔCt).

Quantification of intron 8 retaining (inactive) and normally spliced (active) transcripts of *folylpolyglutamate synthetase* (*FPGS*) was done using qPCR as previously described (30). The ratio was determined by 2^(-ΔCt) where ΔCt is the difference between the Ct values of inactive and active FPGS.

RNA editing of *antizyme inhibitor 1* (*AZIN1*) was determined by a sensitive RNA editing site-specific qPCR as previously described (15) in cell lines and primary mesothelioma cultures established from pleural effusion (31). The Ct value of unedited *AZIN1* is subtracted from that of edited *AZIN1* to obtain a ΔCt and the ratio of edited to unedited *AZIN1* is obtained from 2^(-ΔCt)%.

All primer sequences are listed in Table S3.

### CRISPR/Cas9 knockdown of ADAR2

Human and mouse ADAR2 guide RNA (gRNA) were designed using the CHOPCHOP (https://chopchop.cbu.uib.no/) and GeneAssassin (https://geneassassin.org/) online tools. Three sgRNAs (sequences listed in Table S3) were designed each for human and mouse ADAR2 – 2 within the deaminase domain and 1 in the dsRNA binding domain. ADAR2-targeting gRNAs were cloned into the Cas9-expressing mammalian expression vector pSpCas9(BB)-2A-Puro PX459 (AddGene#158112, #158113, #158114, #158117, #158118, #158119). RN5 (10^5^ cells/w) and Mero95 (1.5× 10^5^ cells/w) cells seeded in 6-well plates were co-transfected with the three plasmids containing the gRNAs (total 400 ng DNA) and Lipofectamine 3000 reagent (Thermo Fisher Scientific) according to manufacturer’s instructions. 48 hours after transfection, cells were selected with puromycin. Individual clones were isolated by seeding cells in 96-well plates at a 1cell/well density. The loss of ADAR2 was determined by protein expression.

### Adar2 cloning, sequencing and transfection

Short isoform of mouse Adar2 cDNA (NM_130895.3) was amplified from RN5 cells using Phusion High Fidelity DNA polymerase and gene specific primers. Adar2 band was cut out and extracted from agarose gel and cloned into the BsmBI site of the BII-ChBtW (AddGene ##175588) vector, which contains an N-terminal Flag-tag and a blasticidine selection marker. The insert was validated by sequencing (done by Microsynth AG, Balgach, Switzerland). All primers are indicated in Table S3. Plasmid containing the short isoform of human ADAR2, pCD3-FLAG-ADAR2S-6xHis, was a kind gift from Prof. Mary A O’Connell (CEITEC Masaryk University). Within this plasmid, ADAR2 contains an N-terminal Flag-tag and C-terminal 6xHis tag, along with a geneticin selection marker (32).

RN5 Adar2 KD (10^5^ cells/w) and Mero95 ADAR2 KD (1.5×10^5^ cells/w) cells were seeded in 6-well plates 24 hours before transfection with ADAR2 expressing plasmids using 400 ng plasmid DNA and Lipofectamine 3000 reagent (Thermo Fisher Scientific) according to manufacturer’s instruction. 48 hours after transfection, RN5 Adar2 KD transfected cells were selected with blasticidine and Mero95 ADAR2 KD cells were selected with geneticin.

### Quantification of Coatomer Protein Complex subunit α (COPA) mRNA editing

PCR to amplify the region flanking the editing site of COPA (Ile164Val) was performed in a total volume of 50μl containing 1x Green Go TaqR Flexi Buffer (Promega), 2 mM MgCl2 solution, 0.2 mM PCR Nucleotide Mix, 0.5 mM of each primer, 1.25 U of Go TaqR G2 Hot Start Polymerase (Promega) and 33.36 ng of cDNA. The primers used are listed in Table S3. Products were confirmed by electrophoresis on a 2% agarose gel and excised. After purification according to the Macherey-Nagel NucleoSpin® Gel and PCR Clean-up protocol, products were sent for Sanger sequencing (done by Microsynth AG, Balgach, Switzerland). Raw sequencing outputs were quantified with Image J software.

The ratio of the area of G peak to the total area of A and G peaks is then reported as a percentage to represent editing levels.

### MTT growth assay

Cells were seeded at 1000 cells/well in 96-well plates. MTT Assay was performed to follow the growth of the cells on days 1, 3 and 7. To perform the MTT assay, medium was aspirated from the wells and cells were incubated in MTT reagent for 90 minutes at 37°C. Then, lysis buffer was added to wells and incubated at 37°C for 2 hours. Absorbance was measured at 570nm using the SpectraMax 340 Microplate reader.

### Cell cycle analysis

Cell cycle analysis was performed with propidium iodide staining. Cells were trypsinised at 80-90% confluence about 48 hours after seeding, and fixed in 70% ethanol overnight. Cells were washed and then treated with RNase (100 µg/mL) in FBS/ PBS for 20 minutes at room temperature before staining with PI (50 µg/mL) at room temperature for 10 minutes on ice. Samples were analyzed using a Attune flow cytometer (Applied Biosystems, Zug, Switzerland) and ModFit LT cell cycle software (Topsham, ME, USA).

### Spheroid formation in vitro and viability assay

To form spheroids, cells (Mero95 WT – 3000 cell/w, Mero95 ADAR2 KD – 8000 cells/w, Mero95 ADAR2 Rescue – 6000 cells/w, RN5 WT – 350 cells/well, RN5 Adar2 KD – 750 cells/well, RN5 Adar2 Rescue – 500 cells/well) were seeded in 96-well ultra-low attachment round-bottomed plates (Sigma-Aldrich). The number of cells was adjusted in order that 4 days after seeding the spheroids reached a diameter of 175-200 μm. Cells were concentrated by gentle centrifugation at 300 x g for 5 min and incubated at 37°C. On day 4 after seeding, the spheroids were treated with folinic acid (Sigma-Aldrich, 47612, 50µM) or pemetrexed (commercial name ‘ALIMTA’, was purchased from Eli Lilly,Vernier/Geneva, Switzerland, 200, 1000 and 5000 µM) or remained untreated for 6 days and viability was analyzed using the CellTiter-Glo Luminescent Cell Viability Assay (Promega, Dübendorf, Switzerland) to determine the adenosine triphosphate (ATP) content. Luminescence was acquired by using a GloMax 96 Microplate Luminometer (Promega). Each experiment was performed in triplicate. For protein and RNA extraction, 20 spheroids per cell line were pooled on day 7 after seeding.

### RNA interference

Silencing of *Mavs* expression in mouse RN5 Adar2 KD cells was done as previously described (16). Silencing of human STING1 or MAVS was done as follows. ON-TARGETplus SMARTpool against *TMEM173, MAVS* and siGENOME nontargeting siRNA pool #2 and DharmaFECT 1 transfection reagent were obtained from Dharmacon. Briefly, siRNA dissolved in siRNA buffer (Dharmacon) was combined with transfection reagent dissolved in OptiMEM (final concentration 0.042%) and incubated for 20 minutes. Then, cells resuspended in normal growth medium were added to the siRNA/DharmaFECT 1 mixture and seeded onto plates, allowing for a final siRNA concentration of 20 nmol/L. Mero95 ADAR2 KD cells (80,000 cells/w in 12-well plate) were plated for whole-cell protein lysates as wells as RNA extraction.

### Senescence-associated β-Gal activity detection

Senescence-associated β-Gal activity was performed as previously described (33, 34). Briefly, 3500 cells/ well were seeded in 48-well plates and grown till 60-70% confluent. After washing with PBS, cells were fixed for 5 minutes in 2% paraformaldehyde, 0.2% glutaraldehyde in PBS. Following one wash with PBS, cells were stained overnight at 37°C in 1mg/ml X-gal, 5mM potassium ferricyanide, 5mM potassium ferrocyanide, 150mM NaCl, 2mM MgCl_2_, 40mM citric acid/ sodium phosphate buffer (pH 6). β-gal positive cells were counted manually from at least 15 different fields of view from each group.

### Intraperitoneal mesothelioma growth and tumor microenvironment

The animal-related experiments were approved by Animal Resources Centre (ARC) University Health Network (UHN). The Animal Use Protocol (AUP) of this study was AUP#6062.9. Exponentially growing RN5 cells (WT, Adar2 KD, Adar2 rescue) were harvested and cell suspension (4×10^6^ cells/200µl PBS) was injected intraperitoneally (i.p.) into C57BL/6 mice. After two weeks, mice were sacrificed to expose the peritoneal cavity. Peritoneal lavage was collected by rinsing with 5 ml of PBS. Lavage was filtered with a cell strainer (Φ70µm) to collect the cells in tumor microenvironment. Retained spheroids were washed with PBS and transferred to a 24-well plate for quantification and image acquisition using an EVOS™ XL Core Cell Imaging System (Thermo Fisher Scientific, Burlington, ON, Canada). Then spheroids were collected in QIAZOL and processed for RNA extraction.

### Flow cytometry

Peritoneal lavage was collected by washing with 5ml PBS per mouse. Lavage single cells prepared by filtering with a cell strainer (40µm in diameter) were used to characterize immune cell populations by flow cytometry. Single cells were pelleted and supernatant was removed and stored at –80°C for cytokine analysis. Flow cytometry analysis was carried out as previously described (15) with the following anti-mouse antibodies: CD45 (clone 30-F11)-PE Cyanine7, CD11b (clone M1/70)-PE Cyanine7, CD68 (LSBio, Seattle, WA)-FITC, F4/80 (Clone: BM8)-APC, F4/80 (Clone: BM8)-PE. All antibodies and isotypes were purchased from eBioscience or BioLegend (San Diego, CA) unless otherwise stated. Becton Dickinson LSR II Flow Cytometer (San Jose, CA) and FACS Diva™ software were used for data acquisition and FlowJo™ software was used for data analysis.

### Measurement of cytokines/chemokines

Peritoneal lavage fluids were concentrated by ultrafiltration through a low-adsorption polyethersulfonate (PES) membrane (mol. mass. cutoff 3kDa, concentrator Pierce PES 3K, Thermofisher). The average concentration factor was 4.2 with a range from 3.1– 5 and it was used to calculate non-concentrated levels. A Bio-Plex mouse cytokine assay (BioRad) for simultaneous quantification of the concentrations of several signaling molecules (Il-6, G-Csf/Csf3, M-Csf/Csf1, Ifn-γ, Ccl2, Cxcl9, Cxcl10) was run according to the recommended procedure. The concentrations were then normalized to the relative CD45^+^CD68^+^F4/80^+^ tumor associated macrophage population, which represents the major source of these cytokines in the peritoneal environment of mesothelioma (15, 35).

### Statistical analysis

The figures represent the mean values from at least three independent experiments. Paired and unpaired t-test, Mann–Whitney, Kruskal Wallis, Fisher’s exact test or one-way as well as two way ANOVA tests or Pearson correlation analysis were used and have been specified when used. To analyze line graphs, area under the curve was determined for each replicate and the data obtained was compared using one-way ANOVA test. Error bars indicate the standard error of the mean. Statistical analysis was performed using Prism 8 (Graphpad 8.0.0).

## Results

### RNA editing in human mesothelioma

In our previous studies, we showed that an increase in A-to-I RNA editing occurs during the development of mesothelioma in a Neurofibromatosis^+/-^ (Nf2^+/-^) mouse model (15, 16). However, the RNA editing landscape in human mesothelioma has not been explored thus far. We therefore used the same genome-wide quantification tool of adenosine-to-inosine editing activity that we had used in the mouse model (16). This computational tool allows computing an index weighting A-to-G mismatches by the number of all adenosines covered in human Alu in RNA-seq data of mesothelial cells (36) and mesothelioma tissue (5, 6). This analysis revealed that A-to-I editing is significantly increased by 1.7-fold in the tumor tissues compared to the normal mesothelial cells (Fig.1A), while C-to-T editing was not only much lower in level compared to A-to-I editing, but there was also no significant difference between normal and tumor tissue. In addition, high editing levels are maintained in mesothelioma primary cultures (Fig. 1A), indicating that it is a feature of tumor cells themselves. Subsequently, we analyzed the nucleotides enriched or underrepresented around editing sites. We observed that the sequence motif (Fig.1B) is consistent with the ADAR-dependent editing signature as we had previously observed in mouse mesothelioma (16), confirming that RNA deamination is dysregulated in human mesothelioma. Furthermore, in both tumor data sets, we found a weak although significant correlation between *ADAR1* and *ADAR2* expression and the level of A-to-I editing as quantified by the A-to-G index (Fig. S1A).

**Figure 1.**
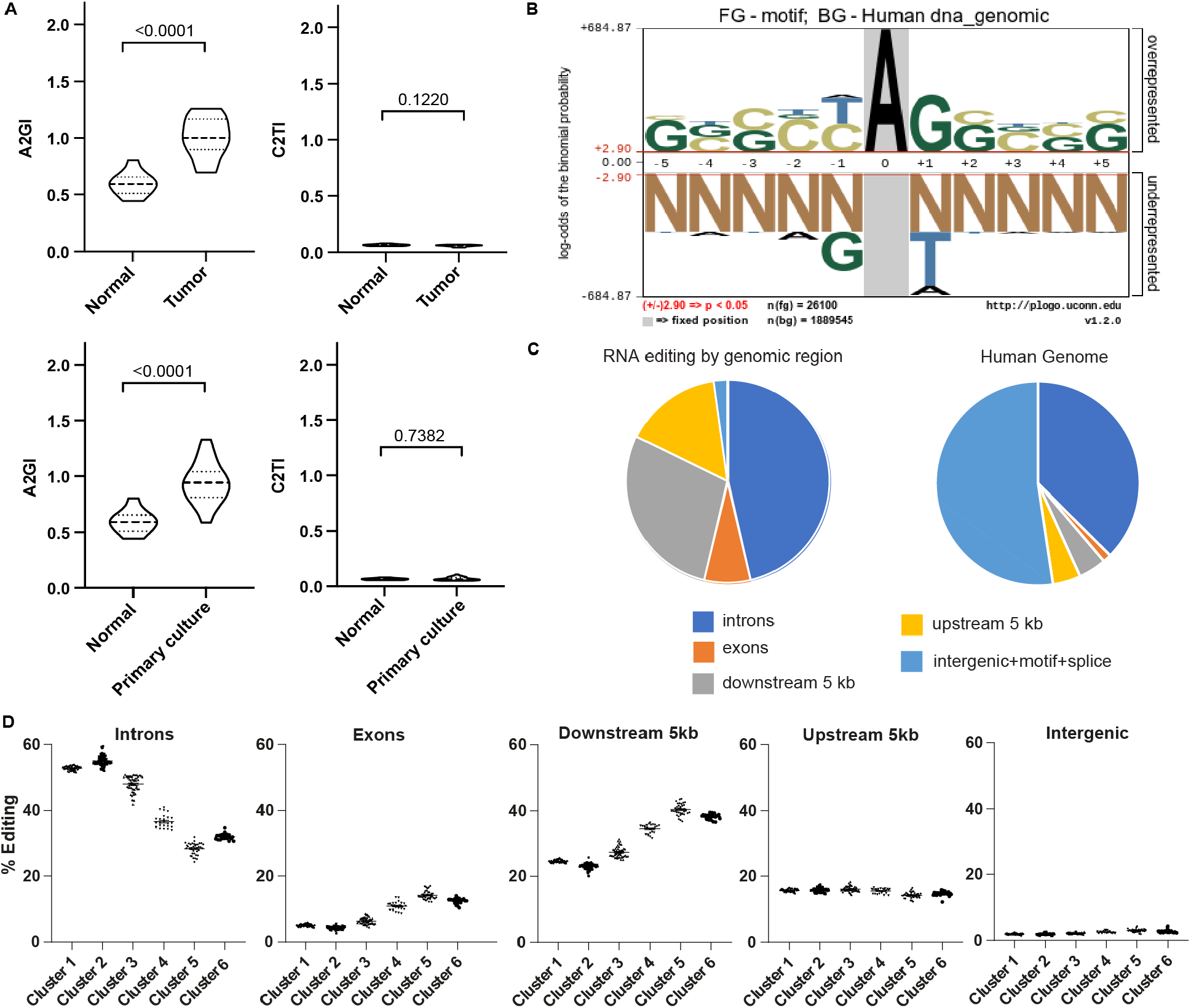
RNA editing in human mesothelioma. **(A)** A to G index (A2GI), which reflects ADAR-dependent RNA editing activity, and C to T index (C2TI) in tumor tissues from TCGA and Bueno’s dataset and primary mesothelioma cultures compared to mesothelial cells. Mann-Whitney test. **(B)** Nucleotides enriched close to editing sites. The height of the nucleotide indicates either the degree of overrepresentation (above the line) or underrepresentation (below the line). **(C)** Genomic localization of RNA editing sites identified in mesothelioma tumor tissue (left) in comparison to the breakdown of genomic regions within the human genome (right). **(D)** Unsupervised clustering of all tumor samples defined 6 groups. Percentage of A-to-I RNA editing within the specified genomic region within each cluster. Significance is shown in Table S4.

To understand the distribution of the identified RNA editing sites, we determined their genomic localization (Fig. 1C). We found that the largest number of editing sites were in the introns, which corresponds to 37% the human genome. However, RNA editing was more than 5-fold enriched in exons and regions 5kb downstream of genes when compared to their relative fraction of the human genome. Unsupervised clustering of the mesothelioma samples based on the genomic localization of RNA editing sites separated them into six groups (Fig.1D, Table S4) with largest editing differences in introns and regions 5kb downstream of genes. Exons include 3’UTR and transcripts (coding sequence) regions (Fig. S1B) and consistent with previous observations (7) the largest fraction of editing occurs in 3’UTR regions. A similar clustering was observed in primary mesothelioma cultures (Fig. S1C, Table S5), suggesting that the editing activity heterogeneity in tumor tissue is also present in tumor cells.

Editing pattern profile in introns is inverted compared to that of exons and 5kb downstream regions, suggesting some dynamics. Increased editing in introns is inversely correlated with pre-mRNA splicing (37) and editing analysis of RNA-seq data of two mesothelioma cell lines treated with a spliceosome inhibitor available from a previous study (5) confirmed that splicing inhibition results in increased editing levels and in dynamic changes between editing in introns and 5kb downstream regions (Fig. S1D).

Altogether, finding different clusters within tumor samples highlight that RNA editing contributes to mesothelioma heterogeneity.

### Expression of ADARs in human mesothelioma and editing heterogeneity association with BAP1

To characterize ADAR expression in mesothelioma cells, we assessed mRNA and protein levels across a panel consisting of 19 human mesothelioma cell lines, one SV-40 immortalized human mesothelial cell line (MeT5A) and primary cells from normal human mesothelium (SDM104, SDM58, and SDM77) (Fig.2A). ADAR1 protein expression was quite homogenous with only 1.5-fold change in expression level between mesothelial and mesothelioma cells (Fig. 2B). On the other hand, ADAR2 protein is heterogeneously expressed in mesothelioma cell lines (Fig.2A), and on an average ADAR2 levels are 2.4-fold higher in mesothelioma when compared to mesothelial cells (Fig.2B). Within this panel of cells, ADAR1 showed no significant correlation between its mRNA and protein while a significant correlation of ADAR2 mRNA expression with its protein levels was observed (Fig. S2A). Nevertheless, we observed a significant correlation between ADAR1, but not ADAR2, mRNA and protein expression and editing of *AZIN1*, a target of both ADAR1 and ADAR2 (38) and one of the 8 recoding sites recently described to increase in cancer (39), in the collection of cell lines (Fig. S2B). The positive correlation between *AZIN1* editing and *ADAR1* mRNA levels was also found in a large panel of primary mesothelioma cultures (Fig. S2C), indicating that *AZIN1* editing most likely reflects ADAR1 abundancy.

**Figure 2.**
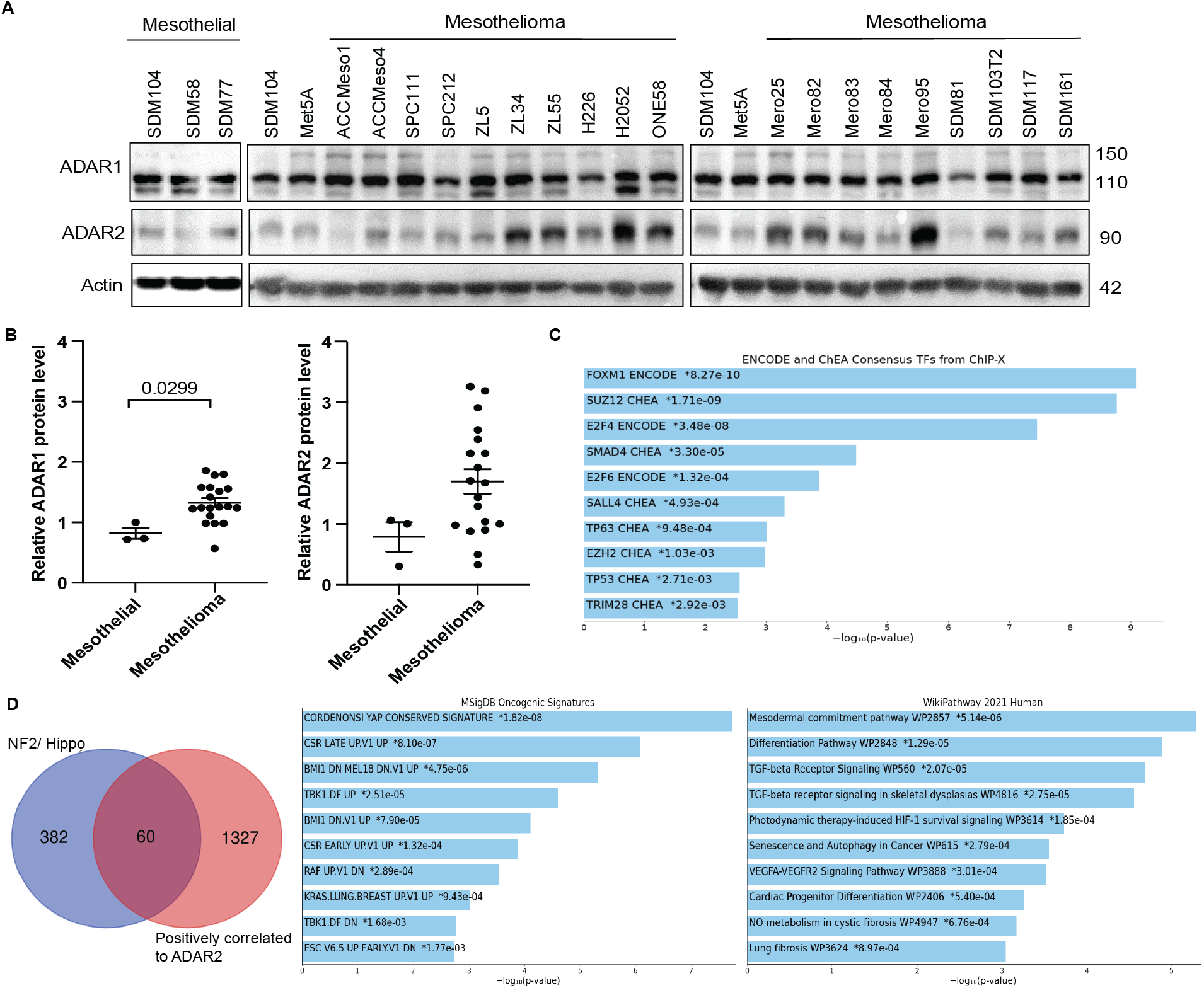
Differential expression of ADAR2 but not of ADAR1 in mesothelioma cells. **(A)** Basal expression of ADAR1 and ADAR2 in 19 mesothelioma cell lines, 3 mesothelial cell lines (SDM104, SDM58, SDM77) and one SV40 immortalized mesothelial cell line (Met5a). Actin was used as the loading control. Size of proteins are represented in kDa. **(B)** ADAR1 and ADAR2 protein quantification in mesothelial and mesothelioma cell lines relative to the SDM104 cell line. Mann-Whitney test. **(C)** Transcription enrichment analysis revealed ENCODE consensus for several transcription factors for genes that are significantly associated with *ADAR2* in mesothelioma TCGA dataset. **(D)** Pathway enrichment analysis for the 60 genes overlapping between *ADAR2* positively correlated genes in mesothelioma TCGA dataset and well characterized genes associated with deregulated NF2/Hippo pathway.

We further analyzed the RNA editing in a set of cell lines based on the expression level of ADAR2 protein. We chose mesothelial cells (SDM104), ADAR2 low cells (ACC Meso1, SPC111, SPC212) and ADAR2 high cells (Mero95, ONE58) and determined RNA editing levels of the codon I164V of *Coatomer Protein Complex subunit α* (*COPA*) mRNA, a specific ADAR2 substrate (40), also part of the 8 differentially edited sites recently described in cancer (39). We found that ADAR2 protein expression was significantly correlated to the A-to-I editing of *COPA* (Fig. S2D).

Since ADAR2 has heterogeneous expression, we aimed at understanding the transcriptional profile in the context of ADAR2 expression in mesothelioma. We therefore extracted genes significantly positively correlated with *ADAR2* in the TCGA dataset using cBioportal (cBioportal.org). We retrieved 1387 (Table S6) genes and transcription enrichment analysis, which focuses on binding sites for transcription factors, revealed ENCODE consensus for Forkhead Box M1 (FOXM1) (Fig. 2C), which is consistent with *ADAR2* being downregulated by YAP silencing (17), since FOXM1 levels are increased by YAP activation (17). To further support this hypothesis, we investigated the overlap between the list of genes significantly positively correlated with *ADAR2*, and well characterized YAP-TEA domain family member 1 (TEAD) genes associated with deregulated NF2/Hippo pathway (41-43). The 60 genes common to both series (Table S7) were not surprisingly resulting in enrichment for YAP conserved signature but also for mesodermal commitment pathway (Fig. 2D).

We consequently concentrated on investigating ADAR2 for several reasons. Firstly, *Adar2* expression was specifically increased in mesothelioma tumor in the experimental mouse model mentioned previously (15), second, ADAR2 levels are higher in mesothelioma compared to mesothelial cells in some mesothelioma cell lines, third, ADAR2 expression is positively controlled by YAP activation and fourth, ADAR2 silencing has been previously observed to inhibit the growth of TTC-MESO1 mesothelioma cells (18).

ADAR2 is localized in the nucleus (8), we therefore analyzed ADAR2 nuclear immunoreactivity in a TMA including 178 mesothelioma tumors diagnostic samples where 50% of the samples were ADAR2 positive (Fig. 3A). ADAR2 positivity was not correlated with histotype.

**Figure 3.**
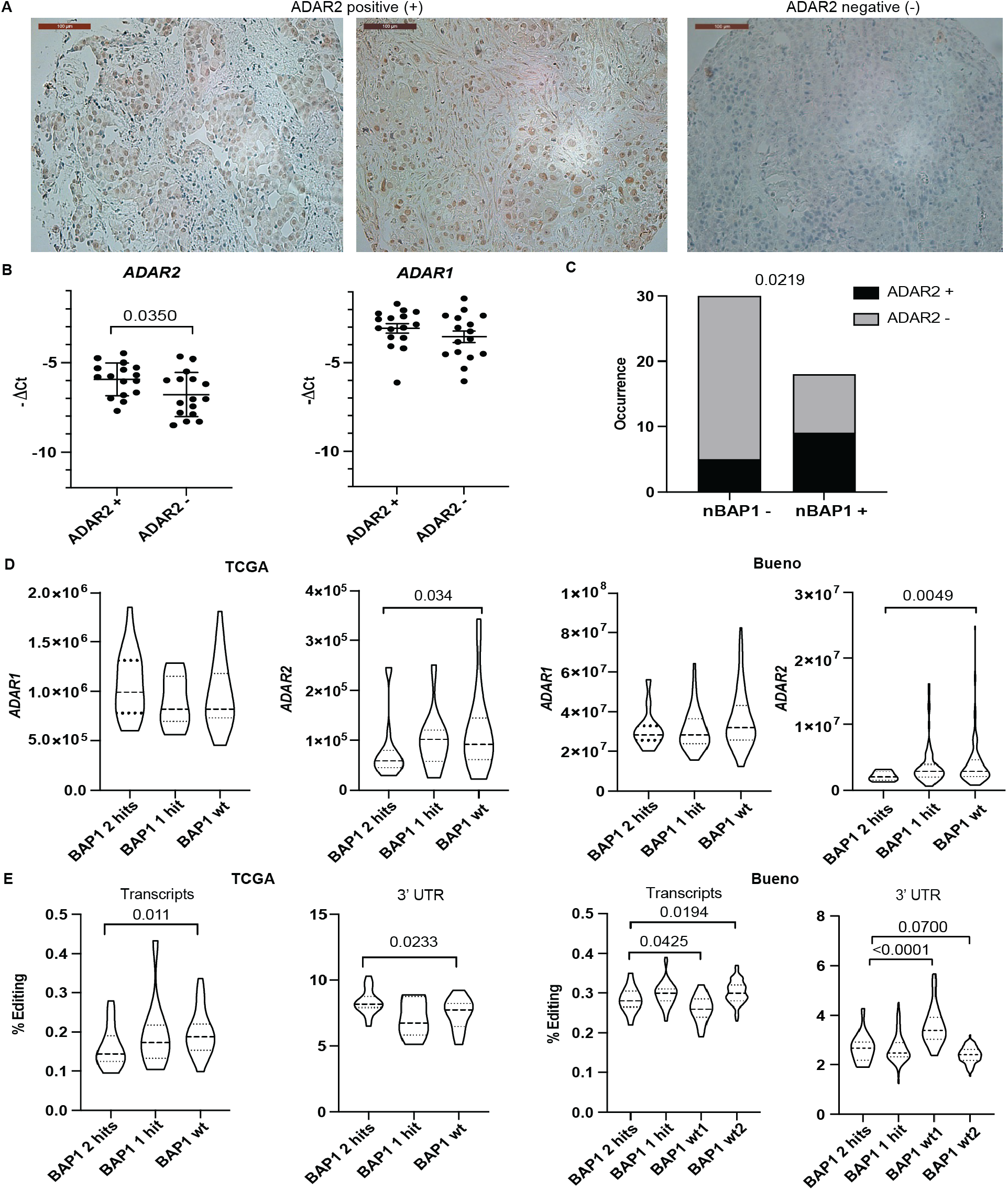
ADAR2 expression is associated with wild-type BAP1. **(A)** Representative images of ADAR2 positive and negative tissues from the tumor microarray. Scale bars are 100μm. **(B)** Quantitative RT-PCR analysis of *ADAR2* and *ADAR1* expression in ADAR2 positive and ADAR2 negative samples. Unpaired t-test. **(C)** Comparison of nuclear BAP1 staining to ADAR2 staining. Two-sided Fisher’s exact test. **(D)** *ADAR1* and *ADAR2* expression in TCGA and Bueno datasets based on the status of BAP1 (wt-wild type BAP1, 1 hit or 2 hit – mutated BAP1) Mann-Whitney test. **(E)** A-to-I RNA editing levels in transcripts and 3’ UTR genomic regions from TCGA and Bueno datasets according to *BAP1* status. Mann-Whitney test.

For a subset of TMA samples (n=33), matching RNA was available and this allowed confirming higher mRNA levels of *ADAR2* in ADAR2 positive samples (Fig.3B), consistent with the correlation between mRNA and protein levels observed in the cell lines (Fig.S2B). No difference in ADAR1 mRNA levels was observed in the ADAR2 negative tissues compared to the ADAR2 positive tissues. No survival difference was observed between nuclear ADAR2 positive vs. negative patients (Fig. S3A).

The destabilization of dsRNA structures upon ADAR activity suppresses the cellular response to dsRNA by preventing type-1 IFN signaling, and the TCGA study (6) has revealed that in mesothelioma this activation occurs in tumors with mutated BAP1. We therefore investigated whether there is an association between ADAR2 and BAP1. For 48 TMA samples, data about nuclear BAP1 immunostaining, which is used as a robust surrogate for wt function (1), and wt BAP1 genotype was also available from a previous study (28) and we observed that ADAR2 positivity is significantly associated with wt BAP1 (Fig.3C). Importantly, the association of *ADAR2*, but not of *ADAR1*, expression with BAP1 status was confirmed in the Bueno (5) and TCGA (6)RNA-seq datasets (Fig. 3D) and was further confirmed in another cohort dataset (CIT series) (24, 44) (Fig. S3B). This was accompanied by a difference in editing in transcripts and 3’UTR according to BAP1 status (Fig. 3E) in the two RNA-seq datasets, while other differences were found only in single datasets (Fig. S3C).

### Characterizing ADAR2 knockdown and rescue cell lines

To investigate the function of ADAR2 in mesothelioma, we used CRISPR/Cas9 technology with three gRNAs to knockdown ADAR2 in the human Mero95 and mouse RN5 cell lines. The RN5 cell line stands as a good model since it expresses both Adar1 and Adar2. Mero95 was chosen from the array of cell lines since it was one of the cell lines having very high expression of ADAR2. The functional domains of ADAR2 include the catalytic deaminase domain and the RNA binding domains. To disrupt both the RNA-binding and catalytic ability of ADAR2, we designed three gRNAs – two within the catalytic domain and one in the RNA-binding domain. We confirmed the decreased levels of protein and mRNA levels (Fig. 4A). Once ADAR2-deficient cell lines were established, we transfected mouse and human Flag-tagged ADAR2 in the ADAR2-knockdown (KD) RN5 and Mero95 cells respectively to generate stable rescue cell lines. The *Adar2* cDNA to rescue Adar2 expression in RN5 Adar2 KD cells was cloned from wild-type RN5 cells where interestingly only the short isoform lacking 10-amino-acid in the deaminase domains (45) is present. The human *ADAR2* to rescue ADAR2 expression in Mero95 cells, corresponded to the equivalent alternative splicing isoform which has been shown to have two-fold higher catalytic activity compared to the longer isoform (46-48). We confirmed the expression of ADAR2 and Flag-tag in the rescue cells (Fig. 4A).

**Figure 4.**
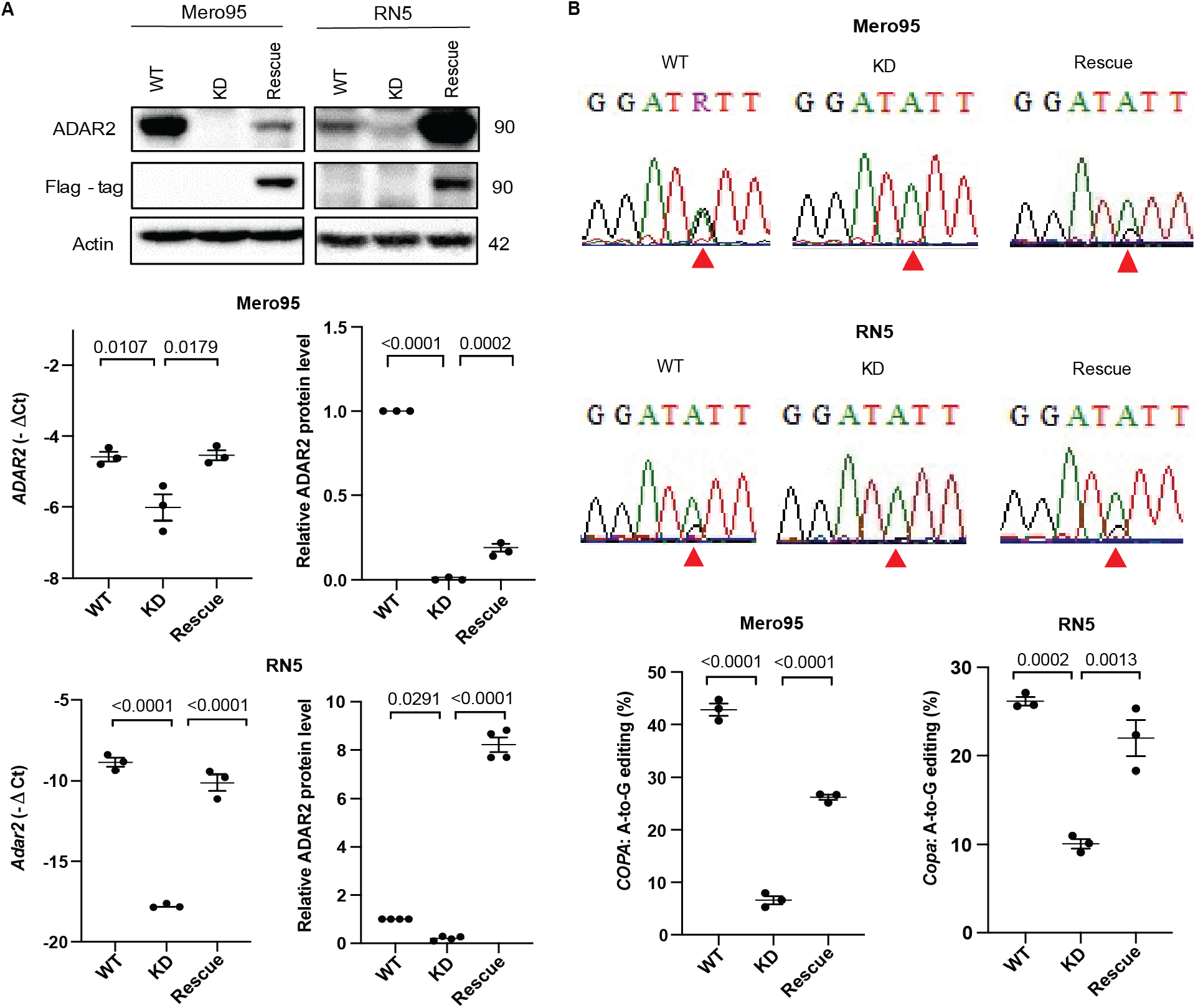
*COPA* editing decreases in ADAR2 KD and is rescued by ADAR2 expression. **(A)** Characterization of ADAR2 protein (size represented in kDa, levels are represented relative to respective WT) and mRNA expression in Mero95 and RN5 ADAR2 knockdown (KD) and ADAR2 rescue cells. One-Way ANOVA with post-hoc Tukey’s test. **(B)** Representative sequence chromatograms of the *COPA* transcript in Mero95 and RN5 WT, KD and rescue cell lines. The sequence surrounding the edited site is conserved in both species. The red arrow indicates the Ile/Val editing position. Quantification of A-to-G changes in *COPA* cDNA from the sequence chromatograms. One-Way ANOVA with multiple post-hoc Tukey’s test.

The efficiency of knockdown and rescue was verified by investigating editing of *COPA* mRNA (Fig.4B). The *COPA* editing in Mero95 ADAR2 KD cells reduced significantly from about 50% to less than 10% and increased to about 30% with the ADAR2-rescue. *COPA* editing in RN5 ADAR2 KD cells reduced significantly but to a lesser extent, which is in accordance with the level of reduction in ADAR2 protein expression in these cells. The ADAR2 rescue in the RN5 cells brought the editing back up almost to the level of the wild-type cells, while no increase in *COPA* editing was observed in mock transfected and selected cells (Fig.S4A). To note, the expression of *COPA* at the RNA level did not change between the three groups in RN5 and Mero95 cells (Fig. S4B).

### ADAR2 deficiency decreases cell growth

To investigate whether different levels of ADAR2 would affect cell growth, cell proliferation using an MTT cell growth assay was examined (Fig.5A). We found that loss of ADAR2 significantly decreased growth of both Mero95 and RN5 cells, and re-expression of ADAR2 was able to significantly rescue cell growth. Cell cycle analysis showed that ADAR2 KD in both Mero95 and RN5 cells resulted in cells accumulating in the G1-phase with a simultaneous decrease in cells within the S-phase, which was rescued by re-expression of ADAR2 (Fig.5B, Table S8).

**Figure 5.**
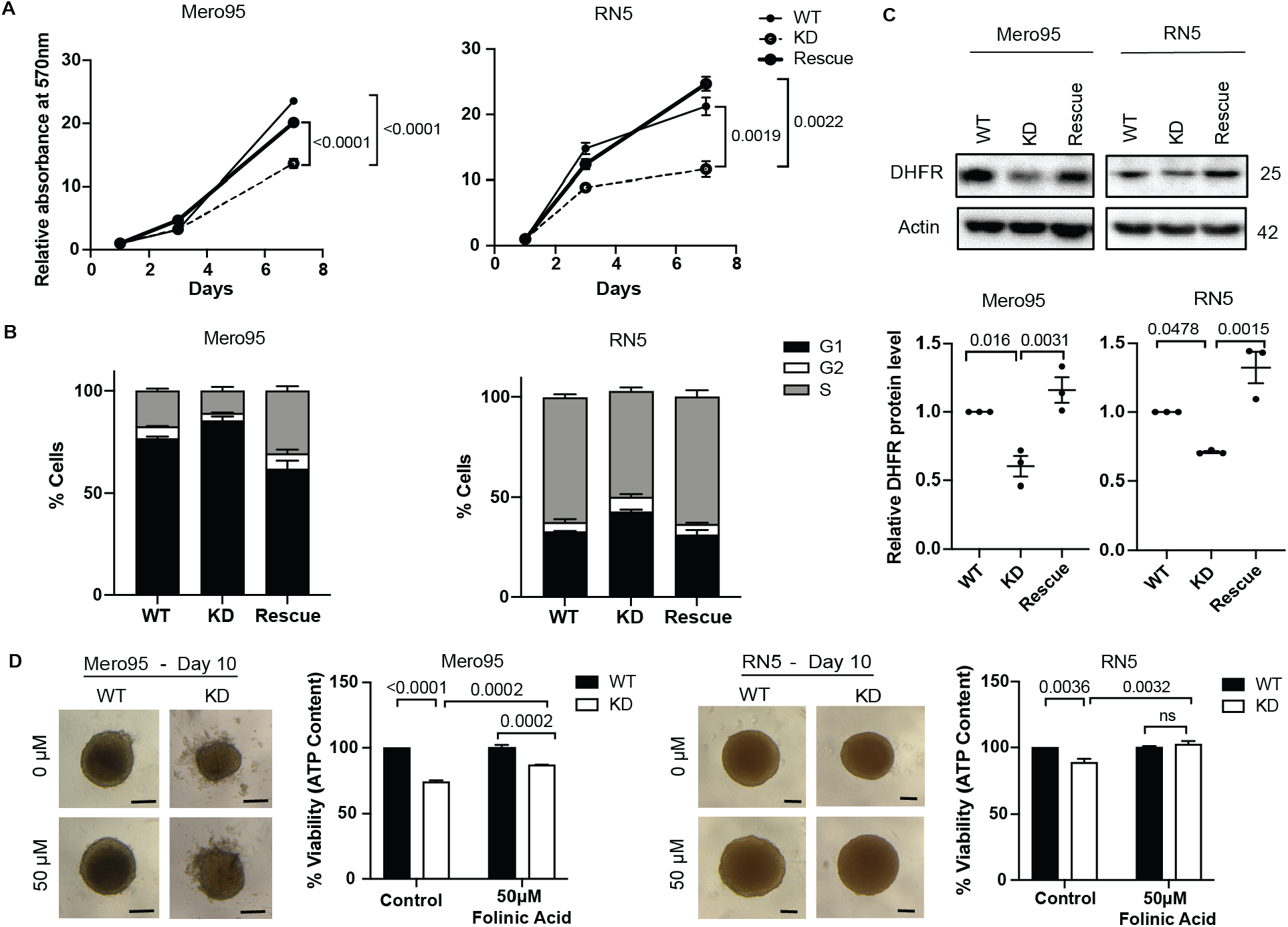
ADAR2 deficiency decreases cell growth. **(A)** MTT assay to assess the growth of Mero95 and RN5 WT, KD and rescue cell lines. Data is relative to the absorbance of respective cell line on Day 1. One-way ANOVA with post-hoc Tukey’s test. **(B)** Cell cycle analysis of Mero95 and RN5 WT, KD and rescue cells. Significance is provided in Table S8. **(C)** DHFR protein expression in Mero95 and RN5 WT, KD and rescue cells. Protein level is represented relative to the respective WT. Size of protein is represented in kDa. One-way ANOVA with post-hoc Tukey’s test **(D)** Representative images of spheroids grown in the presence or absence of folinic acid (50µM) for 6 days. Scale bars are 100µm. Viability of spheroids was quantified using Cell-Titer Glo, which measures ATP content. Two-way ANOVA, with post-hoc Tukey’s test.

One of the targets of ADAR-mediated editing is *dihydrofolate reductase* (*DHFR*) whose expression is regulated by editing in its 3’UTR (49, 50) and high editing levels are present in mesothelioma (Table S9). DHFR plays an important role in cell growth and proliferation by converting dihydrofolates to tetrahydrofolates, required for *de novo* purine and thymidylate synthesis. With loss of ADAR2 we found that DHFR expression decreased in both Mero95 and RN5 cells while its expression increased in ADAR2 rescue cells (Fig. 5C). To validate the role of DHFR in growth control of these cells, we treated the wild type and ADAR2 KD spheroids with folinic acid (49, 51). Folinic acid is a synthetic folate, which is readily converted to tetrahydrofolates even in the absence of enzymes like DHFR (52). Although the cells were seeded in such a way to obtain similar size spheroids on day 4 (day of treatment), by day 10 (day of read-out) the spheroids formed by the ADAR2 KD cells of both RN5 and Mero95 cells were smaller than those formed by the wild-type cells. With the treatment of folinic acid (Fig. 5D), although the rescue in proliferation was only partial, we still observed significant increase in proliferation of the ADAR2 KD cells. This indicates that DHFR indeed plays a role in proliferation of the RN5 and Mero95 cells.

### ADAR2 deficiency sensitizes mesothelioma cells to pemetrexed

Pemetrexed is an antifolate used as a first-line chemotherapy for mesothelioma patients (1). Pemetrexed functions by inhibiting the activity of key folate enzymes such as DHFR and thymidylate synthase (TS) (53). The inhibition of folate metabolism by pemetrexed contributes to ineffective DNA synthesis resulting in failure of tumor cells growth (54). We investigated whether loss of ADAR2 affected the sensitivity of RN5 and Mero95 cells to pemetrexed treatment *in vitro*. High pemetrexed doses are necessary in 3D mesothelioma models compared to the ones usually used in 2D (55), but PM spheroids better represent biological complexity of the tumor *in vivo* (56, 57). Upon pemetrexed treatment, size and cell viability RN5 ADAR2 KD spheroids were reduced (Fig.6A). The Mero95 ADAR2 KD spheroids did not differ much in size to their respective controls, but the debris around the spheroid increased in the ADAR2 KD spheroids compared to wild type. The cell viability assay showed a significant increase in sensitivity to the inhibitory effects of pemetrexed in ADAR2 KD cells. ADAR2 rescue was able to significantly reduce the sensitivity of the cells to pemetrexed treatment (Fig.6A).

**Figure 6.**
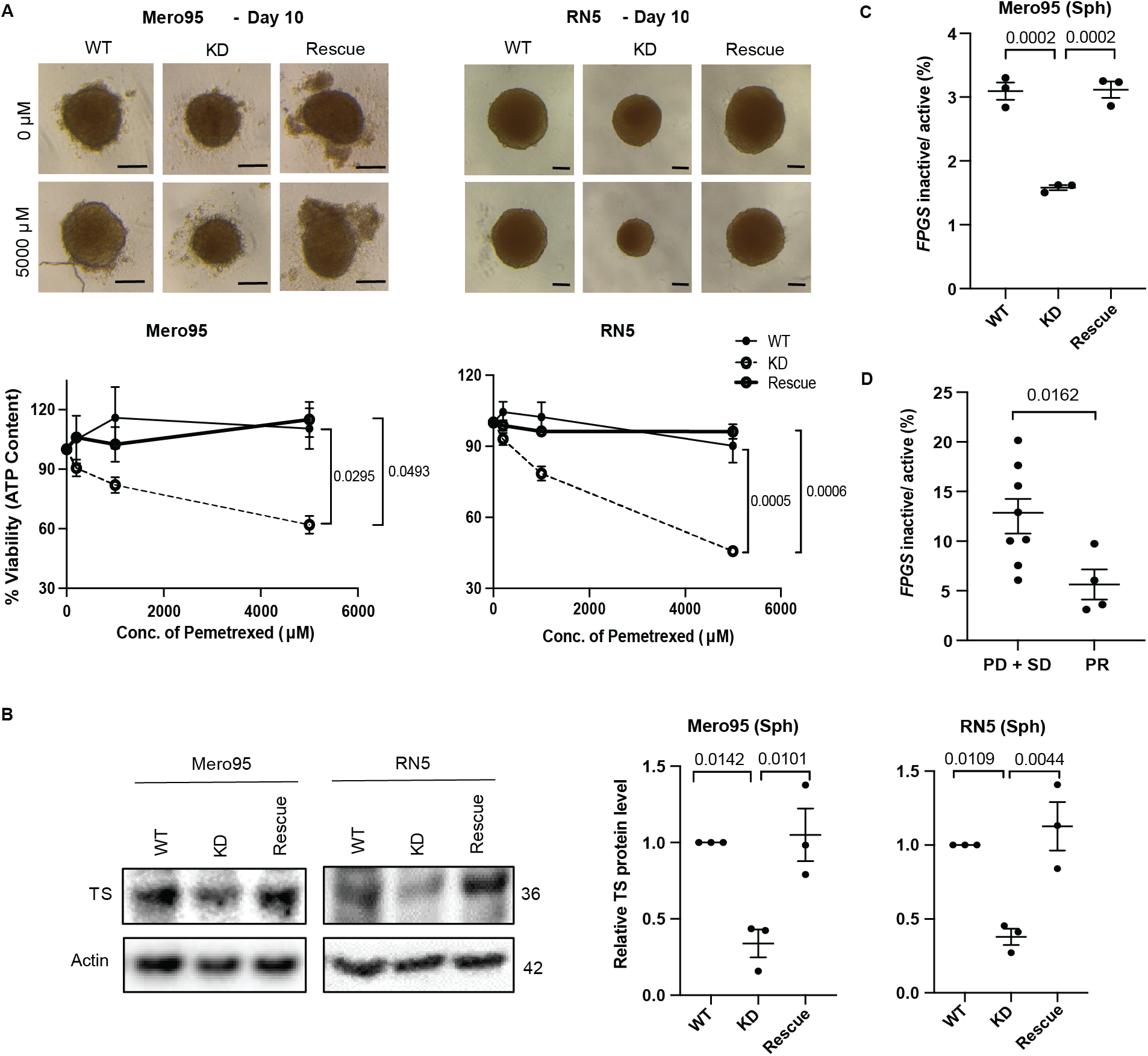
ADAR2 deficiency sensitizes mesothelioma cells to pemetrexed. **(A)** Representative spheroids control or treated with pemetrexed (5000µM) for 6 days. Scale bar is 100µm. Quantification of viability of spheroids after 6 days of pemetrexed treatment. One-Way ANOVA with post-hoc Tukey’s test **(B)** TS expression in Mero95 and RN5 WT, KD and rescue spheroids. Protein levels are represented relative to the respective WT. Sizes are represented in kDa. One-Way ANOVA with post-hoc Tukey’s test. **(C)** Ratio of inactive to active *FPGS* in Mero95 WT, KD and rescue spheroids. One-Way ANOVA with post-hoc Tukey’s test. **(D)** Ratio of inactive to active *FPGS* in patient tumor samples. Progressive disease (PD), stable disease (SD), partial response (PR). Mann-Whitney test.

One of the mechanisms for cancer cells to develop resistance to antifolates such as methotrexate and pemetrexed, is upregulation of the expression of DHFR and TS (53). We analyzed the expression of TS in spheroids and found, like for DHFR, a decrease of its expression in ADAR2 KD cells compared to wild-type cells, which increased again in the ADAR2 rescue cells (Fig.6B).

A recent study has described that altered *folylpolyglutamate synthetase* (*FPGS*) pre-mRNA splicing is associated with unresponsiveness to antifolate agent methotrexate in rheumatoid arthritis (30). FPGS activity adds polyglutamate to antifolate drugs and increases their retention in cells. High levels of intron-retaining *FPGS* transcript leads to reduced activity, hence resistance (30). Because of the dynamic between RNA editing and alternative splicing (37), we analyzed the ratio of the intron 8 retaining (inactive) transcript *vs*. normally spliced (active) FPGS and observed (Fig.6C) that the ratio decreases in Mero95 ADAR2 KD spheroids, which may be part of the sensitizing mechanism. The phenotype was reversed in the ADAR2 rescue spheroids.

We then explored whether ADAR2 expression is associated with chemotherapy response in the subset of patients (n=99), which were treated with pemetrexed/cisplatin induction chemotherapy before surgery. Only a trend for ADAR2 status and response to treatment could be observed (Fig. S5). However, using the RNA, which was available for a subset of patients (n=12), we determined that the ratio of *FPGS* intron 8 retaining transcript *vs*. normally spliced *FPGS* was significantly lower in responders (Fig 6D). Altogether this data indicates that expression of ADAR2 modifies response to antifolate treatment.

### ADAR2 and tumor microenvironment

dsRNA is identified and destabilized by ADARs, thereby dampening the type-1 IFN induced inflammatory response (reviewed in(58)). Although in our previous study we had shown that murine mesothelioma cells bear a high basal level of IFN stimulated genes (ISGs) associated with high levels of endogenous retroviruses (16), in the absence of ADAR2 we observed (Fig.7A) an upregulation of the IFN-response with an increase in expression of the ISGs retinoic acid-inducible gene I (RIG-I) and IFN-induced transmembrane protein 1 (IFITM1) in both Mero95 and RN5 cells. The ADAR2 rescue dampened this upregulated type-1 IFN response (Fig.7A). In the murine model, Mitochondrial antiviral-signaling protein (*Mavs)* silencing was sufficient to effectively reduce Rig-I and Ifitm1 at the protein level (Fig.7B). However, in Mero95 KD cells we observed that the most efficient downregulation of ISG was obtained by silencing stimulator of IFN genes (STING)-encoding *TMEM173* compared to *MAVS* (Fig. S6 and Fig. 7C). This is consistent with the recent observations that ADAR2 activity can dissolve DNA:RNA hybrids (14) decreasing the amount of cytosolic DNA able to activate STING. The release of cytosolic DNA was supported by the activation of senescence-associated β-galactosidase activity (59, 60) in Mero95 ADAR2 KD cells (Fig. 7D).

**Figure 7.**
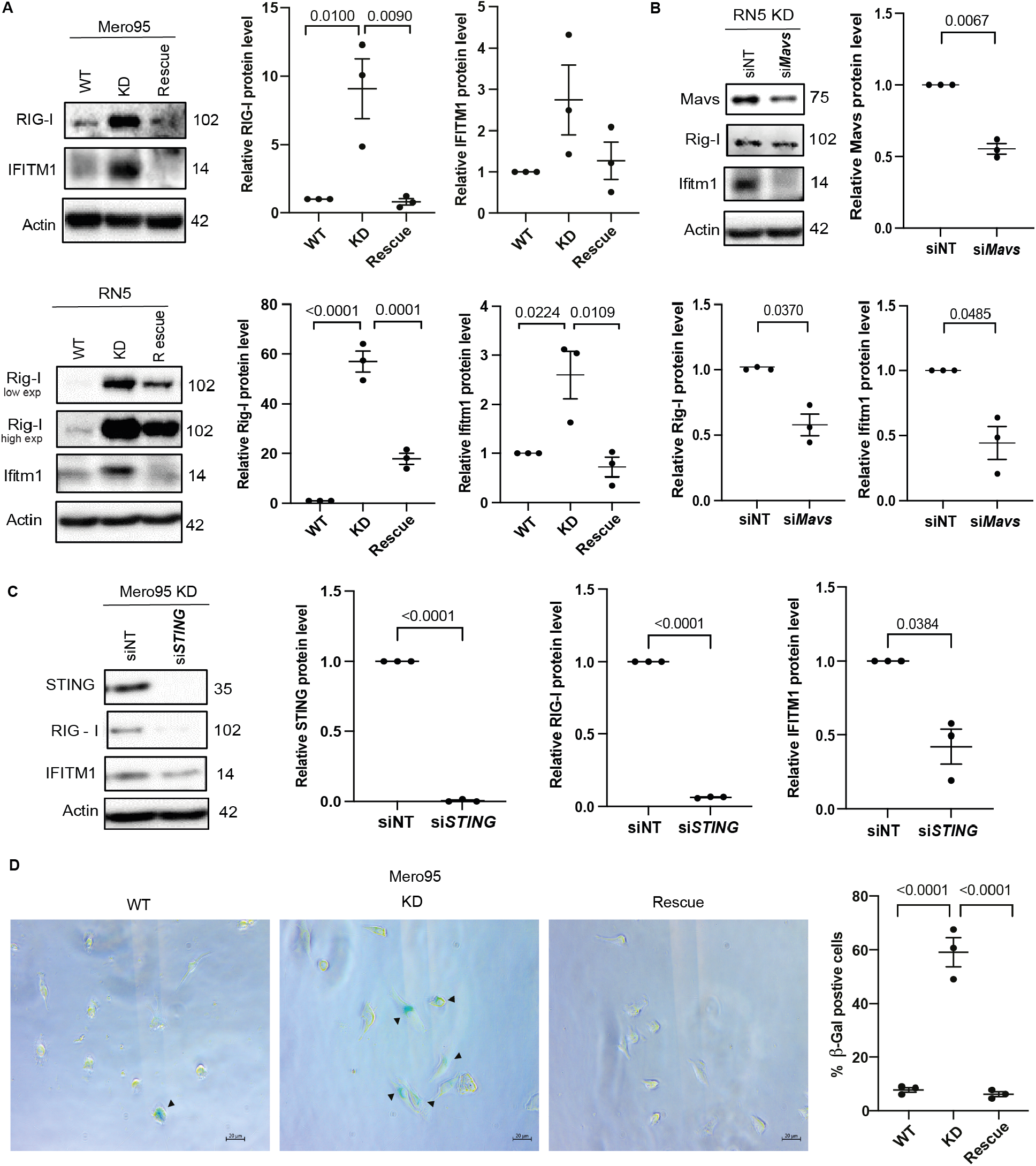
ADAR2 deficiency leads to upregulation of type 1 IFN response. **(A)** Basal expression of ISGs – RIG-I and IFITM1 in Mero95 and RN5 WT KD and rescue cells. Protein levels are represented relative to the respective WT. Protein size is represented in kDa. One-Way ANOVA with post-hoc Tukey’s test. **(B)** Expression of ISGs – Rig-I and Ifitm1 upon silencing of Mavs in RN5 ADAR2 KD cells. Protein levels are represented relative to siNT. Protein size is represented in kDa. Paired t-test. **(C)** Expression of ISGs – RIG-I and IFITM1 upon silencing of STING in Mero95 ADAR2 KD cells. Protein levels are represented relative to siNT. Protein size is represented in kDa. Paired t-test. **(D)** Representative images of senescence-associated β-Gal staining in Mero95 WT, KO and rescue cells (left). Quantification of β-Gal positive cells (right). One-way ANOVA with post-hoc Tukey’s test.

We next aimed at investigating whether Adar2 deficiency alters growth *in vivo*. To that goal, we injected RN5 WT, Adar2 KD, and Adar2 rescue cells into the peritoneal cavity of syngeneic mice, where RN5 proliferation can be followed by counting the number of spheroids formed in the peritoneal lavage (61). Consistent with the observations *in vitro*, Adar2 deficiency resulted in a 12-fold decrease of spheroid number, which however was only slightly rescued in Adar2 expressing cells (Fig. 8A). This was paralleled by a drastic decrease in the number of Cd45^+^Cd68^+^F4/80^+^ tumor associated macrophages (TAM) (Fig. 8B), with a non-significant trend toward an increase with rescue cells. This observation was confirmed using Cd11b^+^ F4/80^+^ macrophage markers (Fig. S7A). Cd11b and F4/80 are canonical macrophage markers and recent studies have identified two sets in peritoneal macrophages characterized by their high (large peritoneal macrophages) *vs*. low (small peritoneal macrophages) expression (62). When we analyzed in the Cd45^+^ and in the Cd11b^+^ series the relative amount of the F4/80_low_ population, we observed an increase of this population in the peritoneal lavage of mice injected with Adar2 deficient cells (Fig. S7A). To further investigate changes in the tumor microenvironment we determined the levels of TAM-produced cyto/chemokines Ccl2, Ifn-γ, Cxcl10, Cxcl9, Csf1, Csf3 and Il-6 in the peritoneal lavage (Fig. 8C and Fig. S7B). Consistent with type-1 IFN activation (63-65), Cxcl10 relative levels were significantly increased and were paralleled by an increase in Ifn-γ in Adar2 KD, while a tendency toward a decrease was observed in the peritoneal lavages from mice injected with Adar2 rescue cells. Ccl2 levels were significantly different in the three groups as well, while for the rest of the investigated cyto/chemokines no significant differences were observed (Fig. S7B). Spheroids forming in the peritoneal cavity were collected and RNA was extracted to investigate mesothelioma and hematopoietic markers. Indeed, TAM are known to associate with tumor cells intraperitoneally injected in mice (35). We compared the expression of mesothelioma markers between *in vitro* and *in vivo* spheroids (Fig. S7C). For the *in vivo* spheroid analysis we added *Ptprc*, which encodes for the Cd45 antigen, to estimate the content of hematopoietic lineage cells. Compared to clustering of *in vitro* spheroids (Fig. S7C, left panel), clustering of *in vivo* spheroids was affected by the content of hematopoietic lineage cells (Fig. S7C, right panel). We therefore explored the expression of a curated gene list of hematopoietic genes involved in lineage differentiation. Unsupervised clustering revealed that the inflammatory profile of Adar2 KD-derived spheroids is distinct from the profile of wild type and rescue-derived spheroids (Fig. 8D), consistent with changes in the chemo/cytokine profile.

**Figure 8.**
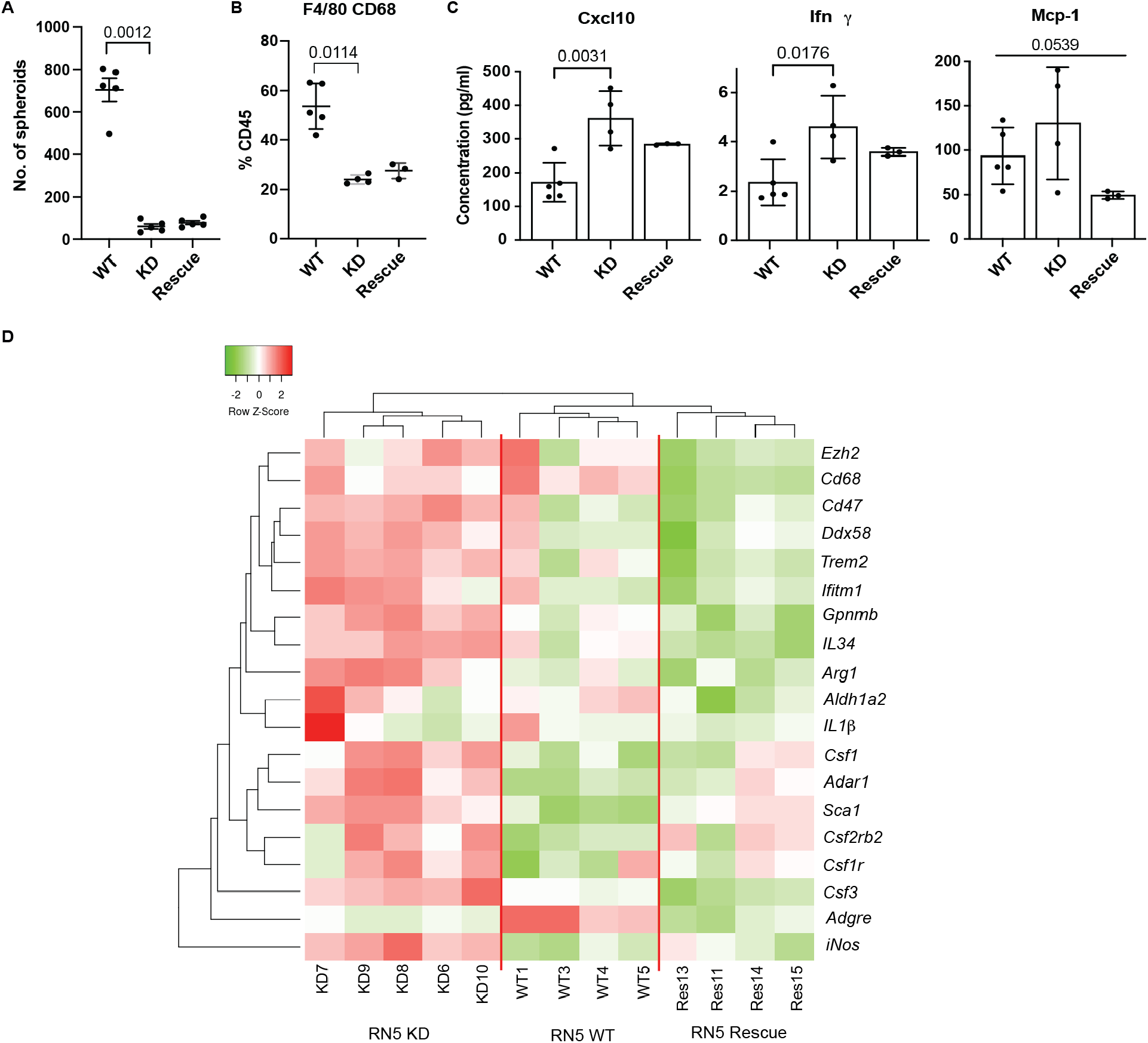
ADAR2 deficiency decreases tumor growth *in vivo* and leads to changes in the tumor microenvironment. RN5 WT, KD and rescue cells were injected i.p. into syngeneic mice and two weeks later the animals were sacrificed and peritoneal lavage was collected to characterize **(A)** spheroids number, one-way ANOVA with post-hoc Tukey’s test, **(B)** Cd45^+^Cd68^+^F4/80^+^ population, Kruskal-Wallis test, **(C)** cytokines/chemokines, one-way ANOVA with post-hoc Tukey’s test. **(D)** Heat map of differentially expressed hematopoietic genes (normalized to *Ptprc* encoding for Cd45) in *in vivo* spheroids. The color bar above indicate upregulation (red) and downregulation (green).

Altogether this data supports the concept that activation of type-1 IFN signaling due to Adar2 deficiency affects the tumor microenvironment.

## Discussion

In this study, we show that ADAR-dependent RNA editing increases in human mesothelioma similarly to what we had observed in the experimental model of mesothelioma development in mice exposed to crocidolite (15, 16). The most abundant editing in mesothelioma was observed in introns, according to what has been described previously (66). The abundant editing in introns is important for the regulation of alternative splicing (37), since both these processes occur co-transcriptionally. Therefore, the identification of clusters with different RNA editing in introns suggests the presence of different splicing in these clusters. Splice variants have been identified in mesothelioma with mutated SF3B1 splicing factor (5), but have not yet been systematically analyzed, although we have documented their occurrence in several mesothelioma relevant genes such as *lncRNA GAS5, CALB2* and *RBM8A* (26, 67, 68) or in major mesothelioma tumor suppressors such as *NF2* (69) and *BAP1*(70). Splice variants have functional consequences including in response to therapy. For instance mesothelioma cells with higher levels of *BAP1⊿* splice variant are more sensitive to poly(ADP-ribose) polymerase inhibition by olaparib, indicating functional consequences of altered splicing (70).

When we consider the rate of editing in specific genomic locations, the most enriched region is the 5 kb downstream of gene as it has also been observed previously (71). Stress-induced transcripts downstream of protein-coding genes have been recently described to significantly contribute to human transcriptome (72) and to maintain nuclear integrity. A recent study investigating RNA editing recoding in human tissues has demonstrated the presence of putative editing complementary sequences up to 15 kb downstream to canonical transcripts, suggesting the existence of extended UTR not included in the canonical RefSeq transcripts (39). The profile of editing in different clusters in the region 5 kb downstream of genes is similar to the editing profile of exons. In the exons, 3’UTR is the most edited region consistent with previous observations (66) and we have recently shown that editing of the 3’UTR of *RBM8A* in mesothelioma cells increases protein levels by counteracting the negative regulation by Musashi2 (26). The opposite profile of clusters in introns and 5 kb downstream region as well as the analysis of RNA-seq data from mesothelioma cell lines treated with a spliceosome inhibitor suggests a dynamic between the two profiles, which needs to be further explored.

We also showed that ADAR-dependent RNA editing varies in some genomic regions according to BAP1 status. Importantly, the expression of ADAR2, but not of ADAR1 is heterogeneous, in both cell lines and clinical samples, and associated with wild-type BAP1. From the point of view of mesothelioma biology, we suggest that ADAR2 expression is part of the landscape of mesothelioma tumors with specific characteristics for two reasons. First, Adar2 is expressed in the potential cell of origin of mesothelioma. Indeed, in a recent single cell transcriptome analysis of twenty mouse organs (73) (https://tabula-muris.ds.czbiohub.org/), single cells from a given organ were identified and sorted using cell surface markers. In the diaphragm, that is often used in studies on mesothelium because this organ is covered by mesothelial tissue on the peritoneal surface, Adar2 is frequently expressed in some mesenchymal stem cells (Sca-1^+^, CD31^-^, CD45^-^). We have described expression of Stem cell antigen 1(Sca-1 also called Ly6A), a gene induced by type-1 IFN (74), in putative mesothelioma stem cells, which are enriched upon therapy in an experimental mouse model (75). The same enrichment of expression of ADAR2 is observed in human mesenchymal stem and some mesothelial cells (https://cells.ucsc.edu/?ds=tabula-sapiens+all&gene=ADARB1). Second, in a experimental animal model of asbestos-induced mesothelioma development (15) we observed a more than two-fold significant increase in expression of *Adar2* in tumors compared to inflamed tissues, but its expression did not differ between sham and inflamed tissue. *Adar2* cDNAs cloned from mouse mesothelioma cells were exclusively derived from short alternatively spliced form, which in human corresponds to the form with the highest catalytic activity. The mechanism of selection of this splice variant is not known and we put forward the hypothesis that tumor cells require high editing capacity, including to dampen type-1 IFN signaling associated with endogenous dsRNA generated by loss of methylation, as we have recently shown (16). Although decrease in type-1 IFN signaling has been most often associated with ADAR1 activity (6, 76-79), our data demonstrate involvement of ADAR2 as well, which is consistent with similar although small effects observed in mice (80). In the murine model, the quenching of the IFN activation by silencing *Mavs* suggests that endogenous dsRNA are the substrate of Adar2. However ADAR2 activity has been recently shown to dissolve DNA:RNA hybrids (14) which form at DNA breaks during transcription and interfere with DNA repair (81). This may result in release of DNA in the cytosol and activation of STING (82). In Mero95 ADAR2 deficient cells, we observed increased levels of the senescence-associated β-galactosidase activity and efficient dampening of type-1 IFN activation by silencing STING-encoding *TMEM173*, indicating an additional involvement of ADAR2 activity in DNA damage response. *In vivo* experiments confirmed that impeding Adar2 activity has a consequence on the tumor microenvironment and this aspect should be further explored namely to better characterize the influence on TAM. Since ADAR2 is associated with wt BAP1, this may help in defining the landscape where mutated BAP1 and low ADAR2 are associated with increased type-1 IFN signaling (6).

Dampening the activation of the innate immune system by ADAR2 activity is part of mesothelioma similar carcinogenesis mechanisms including mutations in the E3 ubiquitin ligase FBXW7 (F-box/WD repeat containing protein) tumor suppressor gene (5, 83-85), which impairs post-transcriptional stabilization of dsRNA sensors (86).

Additional datamining (this manuscript) and experimental observations (17) suggest that ADAR2 expression is downstream of the activation of YAP in mesothelioma cells. YAP is a transcriptional co-activator after interaction in the nucleus with TEAD family of transcription factors (87) resulting in the induction of genes promoting cell proliferation and inhibition of apoptosis (87-90), and we had observed its activation during mesothelioma development in an experimental animal model (15). Hippo cascade regulates YAP via large tumor suppressor homolog 1/2 (LATS1/2)-dependent phosphorylation and subsequent cytosolic sequestration (89, 91, 92). In mesothelioma, due to the NF2 or LATS2 loss, Hippo signaling becomes dysregulated (93-95), although the observation of frequent sub clonal NF2 mutations detected in clinical samples suggests that it may occur later in mesothelioma development (19, 96). Whether mutations in the Hippo pathway correspond to changes in RNA editing should be further explored.

RNA editing maintains DHFR levels, as it has been observed in recent studies (49, 50). As expected as a consequence for decreased levels of the enzyme, which converts dihydrofolate into tetrahydrofolate, a methyl group shuttle required for the de novo synthesis of purines, thymidylic acid and certain amino acids, decreased cell growth was observed upon loss of ADAR2. Pemetrexed and its polyglutamated derivatives inhibit TS, DHFR, and glycinamide ribonucleotide transformylase, all of which are involved in the de novo biosynthesis of thymidine and purine nucleotides. Therefore, pemetrexed treatment induces an imbalance in the cellular nucleotide pool and inhibits nucleic acid biosynthesis, which results in arresting the proliferation of tumor cells. ADAR2 deficient cells expressed lower levels of TS and although intronic TS editing sites (Table S10) are present in RADAR database (97), they have not yet been functionally investigated. However, decreased TS levels might also be a consequence of a slower cell cycle (98). Low TS protein expression and clinical response in pemetrexed/platinum agents treated mesothelioma patients has been controversial (99-101) and no correlation has been observed *in vitro* between TS expression and pemetrexed sensitivity in mesothelioma cell lines (100). However, this has been tested only in 2D conditions. Other mechanisms affecting pemetrexed response include the expression of *FPGS*, although it is also controversial in mesothelioma (99, 100). Novel developments in understanding the mechanisms of sensitization *vs*. resistance to antifolate treatment has revealed the importance of alternative splicing of *FPGS* and although exploratory, due to the low number of available samples, we observed that alternative splicing of *FPGS* changed upon ADAR2 status and was associated with response. This should be further explored but nevertheless provides a hint that ADAR2 activity may be relevant for mesothelioma treatment.

## Conclusions

In conclusion, RNA editing has wide implications on mesothelioma cell growth, response to therapy, and interaction with the microenvironment. In addition, the association of ADAR2 with wt BAP1 is possibly going to help explore some pending questions about BAP1 and type-1 IFN signaling.

### Data accessibility

The datasets supporting the conclusions of this article are available in the Zenodo repository, (10.5281/zenodo.6504941). The TCGA-Mesothelioma cohort was downloaded from the NCBI database of Genotypes and Phenotypes (dbGaP), under phs000178.v10.p8; the Malignant Pleural Mesothelioma cohort from the Bueno study was downloaded from the European Genome-phenome Archive (EGA) under EGAS00001001563 (EGAD00001001915 and EGAD00001001916); the human pluripotent stem cell derived mesothelium was downloaded from the NCBI Gene Expression Omnibus (GEO), under GSE113090 (GSM3096389-GSM3096398)

## Author contributions

*Study concept and design*: EFB, HR, MdP

*Collection, analysis, or interpretation of data*: AH, WQ, LW, MR, MW, JKR, SS, LOG, MS, VSB, CM, CB, JFF, BV, JR, DJ, MdP, EFB

*Clinical records*: IO

*Data Curation Management*: WQ, AH, EFB

*Writing-Drafting of the manuscript*: AH, EFB

*Writing—review & editing*: all authors

*Guidance and scientific input*: HR, WQ, VSB, DJ, MdP, LW, EFB

*Supervision of all aspects of the study*: EFB

*Funding acquisition*: EFB

All authors read and approved the final manuscript.

## Acknowledgements

We thank Dr. O’ Connell for the kind gift of pCD3-FLAG-ADAR2S-6xHis plasmid.

